# Integrated Detoxification, Bioremoval and Ohmic-Heating Recovery of Lead and Cadmium by *Escherichia coli* K-12 MG1655

**DOI:** 10.64898/2025.12.06.692726

**Authors:** Nnabueze Darlington Nnaji, Chukwudi U Anyanwu, Taghi Miri, Helen Onyeaka

**Author notes:** Authors to whom correspondence should be addressed –.

## Abstract

Lead (Pb) and cadmium (Cd) remain among the most persistent and hazardous heavy-metal contaminants in industrial effluents, posing severe risks to ecosystems and human health due to their non-biodegradable nature and high toxicity. In response to the limitations of conventional chemical remediation technologies, this study evaluates the potential of *Escherichia coli* K-12 MG1655 to function as a microbially driven system for the detoxification, sequestration and recovery of Pb and Cd. Emphasis is placed on oxalic acid production as a mechanistic basis for metal tolerance. High-performance liquid chromatography confirmed that *E. coli* K-12 MG1655 synthesises oxalic acid under metal stress, with Pb exposure eliciting the highest oxalate output, providing biochemical evidence for metal–oxalate complexation as a key detoxification strategy. Bioaccumulation studies using inductively coupled plasma–optical emission spectrometry revealed exceptional metal removal efficiencies, reaching 99.94% for Pb and 97.77% for Cd at 1000 ppm, while Pb + Cd mixed-metal systems maintained high overall uptake (98.19%). These results demonstrate that *E. coli* can sequester metals across a wide concentration range with minimal inhibition from competitive ion interactions. Metal recovery from loaded biomass was evaluated through acid desorption and Ohmic heating. Nitric acid (0.1 M HNO_3_) achieved the highest recovery efficiencies (Pb: 98.5%; Cd: 91.5%), whereas Ohmic heating yielded moderate (Pb: 45.38%; Cd: 45.83%) but environmentally favourable recovery without chemical additives. The integrated findings illustrate a complete microbial remediation–recovery cycle encompassing detoxification via oxalic acid, high-efficiency metal sequestration and effective downstream recovery. This integrative study establishes *E. coli* K-12 MG1655 as a promising candidate for closed-loop bioremediation systems linking detoxification, sequestration and recovery of heavy metals.

**Impact statement:** This study addresses a major gap in microbial bioremediation research by integrating the interconnected processes of detoxification, metal bioaccumulation and metal recovery within a single microbial platform. By demonstrating that oxalic-acid-driven detoxification directly enhances bioaccumulation performance and enables subsequent metal release through either dilute acid or ohmic-heating regeneration, this work provides a unified framework linking microbial physiology with practical recovery technologies. The study advances the field by showing how microbial systems can be engineered into circular, regenerable bioprocesses, reducing dependency on chemically intensive methods and offering scalable, sustainable solutions for the remediation of metal-contaminated environments.

## Introduction

Heavy metal contamination continues to pose a serious global threat to environmental and public health, with the persistence and toxicity of these metals contributing to increasing ecological crises across both industrialised and developing regions. According to the World Health Organization [1], heavy-metal pollution is one of the key public-health issues, particularly in low-income countries where poor management of industrial waste often subjects the population to polluted soil and water. Regulatory frameworks such as the European Union Water Framework Directive [2] and U.S. EPA priority pollutant listings [3] underscore the international recognition of Pb and Cd as high-risk toxicants. Pb is considered the most dangerous contaminant because of its neurotoxicity and Cd has severe renal, skeletal and carcinogenic effects and is ranked the sixth on the ATSDR toxicity list [4]. These issues are not far-fetched, empirical measurements in highly industrialised regions like the Niger Delta have found that the concentration of Pb and Cd in residents reaches up to six times of the acceptable limit, further increasing the cumulative human-health burden of chronic exposure [5]. These cases highlight the scale of the contamination problem and support the importance of new approaches which should be sustainable and socio-economically available.

The traditional heavy-metal removal methods, such as chemical precipitation, ion exchange and electrochemical methods, have been widely used but are limited by the high cost of operation, partial recovery of metals, and generation of toxic sludge [6]. These constraints depict the inefficient nature of the remediation measures that are based only on physicochemical principles especially when used in complex effluent matrices or in low concentrations of metal ions. The further increase in industrialisation, urbanisation and mining has also increased anthropogenic metal emissions [7], which exceeds the ability of traditional technologies to provide effective environmental protection. As a result, the focus of research has changed to more sustainable, biologically inspired methods that would be able to tackle ecological and economic limitations. The new paradigm focuses on the resource circularity, low-energy needs, and ecologically friendly metal detoxification and recovery pathways.

Bioremediation has therefore become a trend, which is an environmentally friendly alternative that is based on living organisms, particularly microorganisms that are able to accumulate, transform, or immobilise the toxic metals [8]. *E. coli* has been distinguished in this field due to its wide metabolic flexibility, genetic plasticity and quick adaptation to environmental stresses. Past research shows that the interaction *E. coli* with a wide range of heavy metals can modulate biosorption through cell wall components, transporter systems, and stress-related biochemical pathways [9]. Another factor that demonstrates bacterial metabolic flexibility is the release of organic acids in response to toxic stress, such as oxalic acid, a metabolite that can bind metal ions to form stable, insoluble oxalate complexes [10]. This is commonly known as microbial detoxification, which reduces reactive oxygen species, ensures metal homeostasis, and reduces cellular toxicity [11]. Similar results are reported in parallel studies involving microbial strains that are exposed to Pb, which increases the production of oxalic acid [12], this supports the general hypothesis that organic acid production is an evolutionary response to oxidative and ionic stress.

Although significant progress has been made, the existing literature has a disjointed perception of how the processes of detoxification, metal uptake and biorecovery overlap in a single microbial system. Most existing studies on heavy metal removal focus on either the efficiency of biosorption [13, 14], the metal-tolerance mechanisms [15] or desorption strategies [16] in isolation, with limited attention to how these steps can be integrated into a single, regenerable process. This siloed approach obscures how metabolic responses influence metal sequestration efficiency or how intracellular and extracellular detoxification pathways may determine the feasibility of downstream metal recovery. Although the uptake of heavy metals has been described in systems, little has been done to examine the dynamic nature of the metabolites like oxalic acid in determining the sequestration and recovery potential. In addition, although acid desorption has proven to be an efficient recovery technique [17], other techniques, including Ohmic heating, which is a relatively new technology, are under-researched and not well incorporated into the circular bioremediation framework.

The lack of integrative studies makes several critical questions unanswered. How does *E. coli* coordinate cellular defences including membrane stabilisation, Extracellular Polymeric Substances (EPS) restructuring, enzymatic protection and organic acid secretion in response to metal stress? What is the relationship between oxalic acid production and metal bioaccumulation efficiency under single and mixed metal exposure? Most importantly, what can be done to use these biological processes to enable successful metal biorecovery, and in turn, lessen the use of chemically intensive approaches and enable sustainable circular economies? Addressing these gaps is essential to further the science of microbial bioremediation and to attain scalable, real-world solutions that can ameliorate the heavy metals contamination globally.

Here, we establish an integrated platform based on non-pathogenic *Escherichia coli* K-12 that (i) employs oxalic-acid-mediated detoxification to maintain viability at Pb and Cd concentrations up to 1000 mg/L, (ii) delivers high-efficiency bioremoval of Pb and Cd in single and binary metal systems, and (iii) evaluates both dilute nitric acid and ohmic heating as regeneration routes for metal recovery. To our knowledge, this is the first study to explicitly link oxalate-driven detoxification with ohmic-heating-assisted recovery of Pb and Cd in a unified bacterial system, bridging microbial physiology and process engineering for metal-laden effluents.

Therefore, this study aims to elucidate the detoxification, bioaccumulation as well as biorecovery of Pb and Cd-exposed *Escherichia coli* K-12 MG1655, including oxalic-acid-mediated tolerance and comparative study of acid and ohmic-heating recovery processes. Through a unified examination of microbial physiology, metabolic responses and recovery pathways, this work seeks to advance a more holistic understanding of microbial metal processing and contribute to the development of sustainable, circular bioremediation frameworks.

## Materials and Methods

### Bacterial Culture and Heavy Metal Exposure

A laboratory strain of *Escherichia coli* K-12 MG1655 was obtained from the microbial culture collection of the Biochemistry Laboratory, School of Chemical Engineering, University of Birmingham. The strain was revived by transferring a lyophilised bead into nutrient broth and incubated at 37 °C and 150 rpm within 24 hrs. Purity of culture was established through streak plating on nutrient agar. To achieve physiological homogeneity in the presence of metal, the culture was allowed to reach mid-logarithmic growth and was spectrophotometrically monitored at 600nm with OD_600_ values between 0.5 and 1.0 considered suitable for the experiment. Cultures with a value above this were diluted accordingly.

Analytical-grade lead nitrate [Pb(NO_3_)_2_] and cadmium nitrate tetrahydrate [Cd(NO_3_)_2_·4H_2_O] were used as the sources of metals. Stock solutions were made by dissolving 1.6 g Pb(NO_3_)_2_ and 2.74 g Cd(NO_3_)_2_·4H_2_O in 1 L of deionized water; 100 ppm, 200 ppm, and 500 ppm stock solutions were prepared by diluting the stock solutions. A 0.1 M NaOH or HCl was added to buffer the metal solutions to pH 7.0 and counter the pH-dependent changes in metal bioavailability.

Mineral salts medium (MSM) was composed of 1.73 g/L K_2_HPO_4_, 0.1 g/L MgSO_4_·7H_2_O, 1 g/L NH_4_NO_3_, 0.03 g/L FeSO_4_·7H_2_O and 5 g/L yeast extract. For each assay, 1 mL inoculum was inoculated in a solution of 40 mL MSM, 20 mL sterile glucose solution and 40 mL metal solution. The metal free control consisted of 80 mL MSM and 20 mL glucose solution. Cultures were incubated at 37 °C and agitated in a rotary shaker at 150 rpm for 24 h to achieve uniform aeration and metabolic activity in the presence of metals.

### Quantification of Oxalic Acid

High-performance liquid chromatography (HPLC) was conducted on the 1000 ppm treatment using an Agilent 1260 Infinity II system (Santa Clara, CA, USA) to explain the production of oxalic acid as a possible detoxification product. After 24 h, culture samples treated with Pb and Cd were centrifuged to remove biomass; the supernatants were filtered with 0.22 µm membranes to remove any residual particulates that could affect chromatographic resolution. The standard of oxalic acid at 50 ppm was prepared and analyzed to determine the reference retention time and the corresponding peak area, which defines the calibration baseline required to perform absolute quantification. Reverse-phase C18 columns (250 mm x 4.6 mm, 5 um) were used with flow rates of 1.0-1.2 mL/min ^−1^ and UV detection at 240, 280, 320, and 350nm. The mobile phase mixture was 85 % acetonitrile and 15 % water. The identity of the peaks in chromatograms of experimental samples was verified against the standard; the retention times were used as the main diagnostic criterion. The calibration curve was obtained using the 50 ppm standard, and the areas of the peaks were determined and quantified to obtain the concentration of oxalic acid in each sample.

### Fourier Transform Infrared Characterization of Secreted Oxalic Acid Following 1000 ppm Pb Exposure

FTIR was used to characterize oxalic acid synthesized by *E. coli* K-12 MG1655 exposed to 1000 ppm Pb. After 24 h exposure to metals, centrifugation was done using a Sigma Refrigerated 2023 (1320/G13) centrifuge at 10,000 × g in 20 minutes at room temperature to separate the bacteria cells and take the supernatant, which contained secreted Oxalic acid Fourier transform infrared spectra were recorded with Bruker Vertex 80 FTIR spectrometer with an attenuated total reflectance (ATR) accessory. Acquisition of the spectrums was done at a wavenumber range of 4000 −650 cm^−1^.

### Bioaccumulation Studies

Bioaccumulation of Pb and Cd by *E. coli* was quantified using the Perkin Elmer Optima 8000 inductively coupled plasma–optical emission spectrometry (ICP-OES). Cultures were centrifuged after 24 h exposure to metals to separate supernatant and bacterial pellets. Supernatants were retained for residual metal analysis, and pellets were digested in acid to guarantee that all intracellular and surface-bound metals are released. Concurrent ICP-OES of the two fractions gave accurate measurements of metal uptake. Bioaccumulation efficiency was calculated using the established equation:

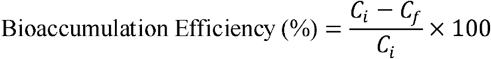

where *C*_*i*_ represents the initial metal concentration and *C*_*f*_ the final concentration in the supernatant. Experiments were conducted in both single-metal and mixed-metal systems to investigate competitive interactions.

### Biorecovery Studies

Biorecovery experiments were used to test the viability of recovering Pb and Cd in metal-contaminated biomass of *E. coli* K12 MG1655 using two different techniques: acid-based desorption and Ohmic heating. This experiment was carried out for the 200 ppm set ups. A 15 mL culture containing *E. coli* biomass from test setup after 24 hours was centrifuged at 10,000 × g for 10 minutes. The pellet was collected and thoroughly washed with PBS to remove any loosely bound heavy metals on the surface. Acid solutions of HCl at 0.01 M, 0.1 M and HNO_3_ at 0.01M and 0.1M, were prepared in sufficient quantities. After decanting the supernatant and leaving the *E. coli* biomass, 15 mL of either 0.01 M or 0.1 M HCl, or 0.01 M or 0.1 M HNO_3_, was added. The biomass was fully immersed in the acid solution and placed in a shaker to gently mix the solution, promoting effective contact between the biomass and the acid. The biomass was incubated in the acid solution for 1 hour at room temperature. After the incubation period, the biomass was separated from the acid solution by filtration. The supernatant, now containing the desorbed heavy metals, was collected in a clean container for ICP-OES analysis.

The second method of biorecovery involved ohmic heating. A resuspension of 50 mL of deionised water of the biomass of *E. coli* was made in a three-necked, 100 mL glass reactor with two stainless-steel electrodes 4.25 cm apart. The ohmic heating system was operated following the procedure described by Chu *et al*. [18]. The setup was operated at 30 V, with the temperature maintained about 30–35°C over a 10-minute period. This created cell disruption and release of metals without the use of chemical reagents, hence minimizing secondary waste. Further filtration provided a supernatant that was subjected to ICP-OES. Recovery efficiency for each method was calculated using the formula:

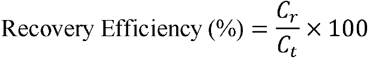

Where *C*_*r*_ is the concentration of recovered metal and *C*_*t*_ the total metal content initially present in the biomass.

### Statistical Analysis

Experiments were conducted in triplicates and the data obtained were subjected to robust statistical treatment to ensure reliability and reproducibility. Analysis of variance (ANOVA) was used to identify significant differences in treatment means with special focus on comparing the bioaccumulation rates and recovery efficiencies under experimental conditions. Where ANOVA identified significant variation, least significant difference (LSD) tests were used to identify pairwise differences at the confidence level of p < 0.05.

## Results

### Oxalic Acid Production Under Metal Stress

Quantification of oxalic acid in cultures of *E. coli* K-12 MG1655 exposed to Pb (1000 ppm) and Cd (1000 ppm) and their combination was determined using HPLC. A 50 ppm standard of oxalic acid was used to compare samples to detect oxalic acid produced by *E. coli* under metal stress based on retention times, peak areas, and peak heights. The known standard had a retention time of 1.273 minutes, with peak area of 1.135 mAUmin, and peak height of 6.313 mAU (Figures 1a). This result served as reference points for the subsequent analysis of *E. coli* samples.

**Figure 1:**
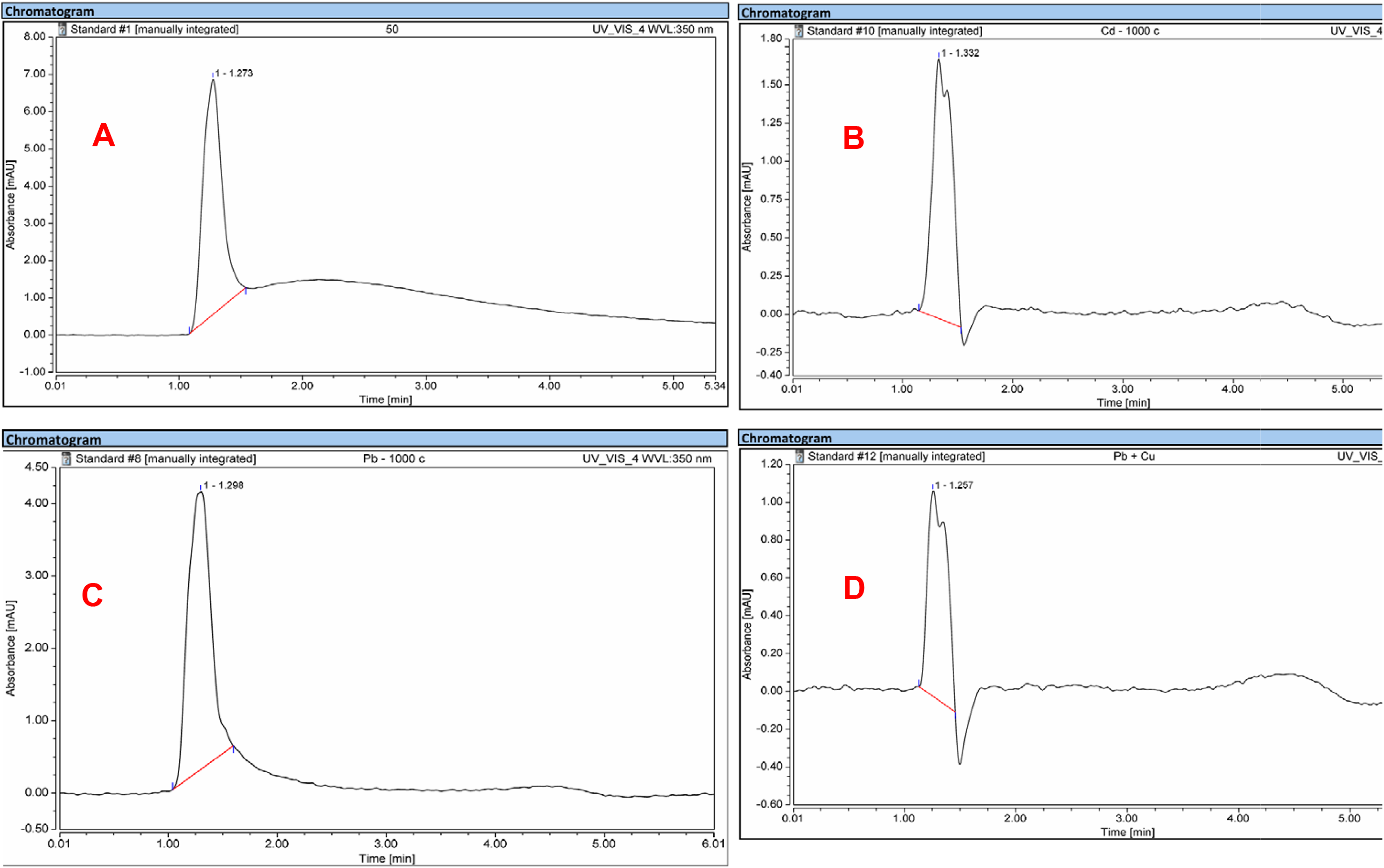
HPLC Chromatogram of - **A:** Known Oxalic Acid Standard (50 ppm); **B:** Oxalic Acid from *E. coli* Exposed to 1000 ppm Pb; **C**: Oxalic Acid f *E. coli* Exposed to 1000 ppm Cd; **D:** Oxalic Acid from *E. coli* Exposed to 1000 ppm Pb + Cd.

The retention time in cultures exposed to 1000 ppm Pb was slightly delayed at 1.298 min and the peak area was 0.937mAUmin and the peak height was 3.831mAU (Figure 1b). On the other hand, the retention time of 1000 ppm Cd was 1.332 min with the peak area being significantly smaller (0.345mAUmin) and the peak height (1.696mAU) being lower (Figure 1c). The culture exposed to the combined stress of Pb and Cd (1000ppm each) had the shortest retention time of 1.257min, the lowest peak area of 0.217mAUmin and a peak height of 1.086mAU (Figure 1d). These findings prove that *E. coli* K-12 MG1655 produces measurable levels of oxalic acid during metal stress, and the levels of production differ depending on the type of metal exposure. The highest oxalic acid production was observed in the 1000 ppm Pb condition as indicated by the larger peak area and height compared to 1000 ppm Cd or the 1000 ppm Pb + Cd exposure, indicating that Pb stress could cause a more significant metabolic response which includes oxalic acid production *in E. coli*.

### FTIR Evidence of Oxalic Acid Synthesis

FTIR analysis of the culture supernatant showed clear spectral differences between Pb-treated samples and the control (Figure 2). The control supernatant exhibited strong O–H/N–H stretching (3359 cm^−1^), carboxylate vibrations (1421 cm^−1^), and multiple bands between 1200–1000 cm^−1^ attributed to extracellular metabolites. In contrast, Pb-exposed supernatant showed markedly reduced intensities across these organic functional groups, consistent with oxalic-acid consumption during metal complexation. A distinct band at 1033 cm^−1^ appeared exclusively in the Pb-treated supernatant, corresponding to C–O–Pb vibrations characteristic of oxalate–metal species. These findings indicate that extracellular oxalic acid synthesized by *E. coli* reacted with Pb^2+^ to form Pb-oxalate complexes detectable in the supernatant.

**Figure 2.**
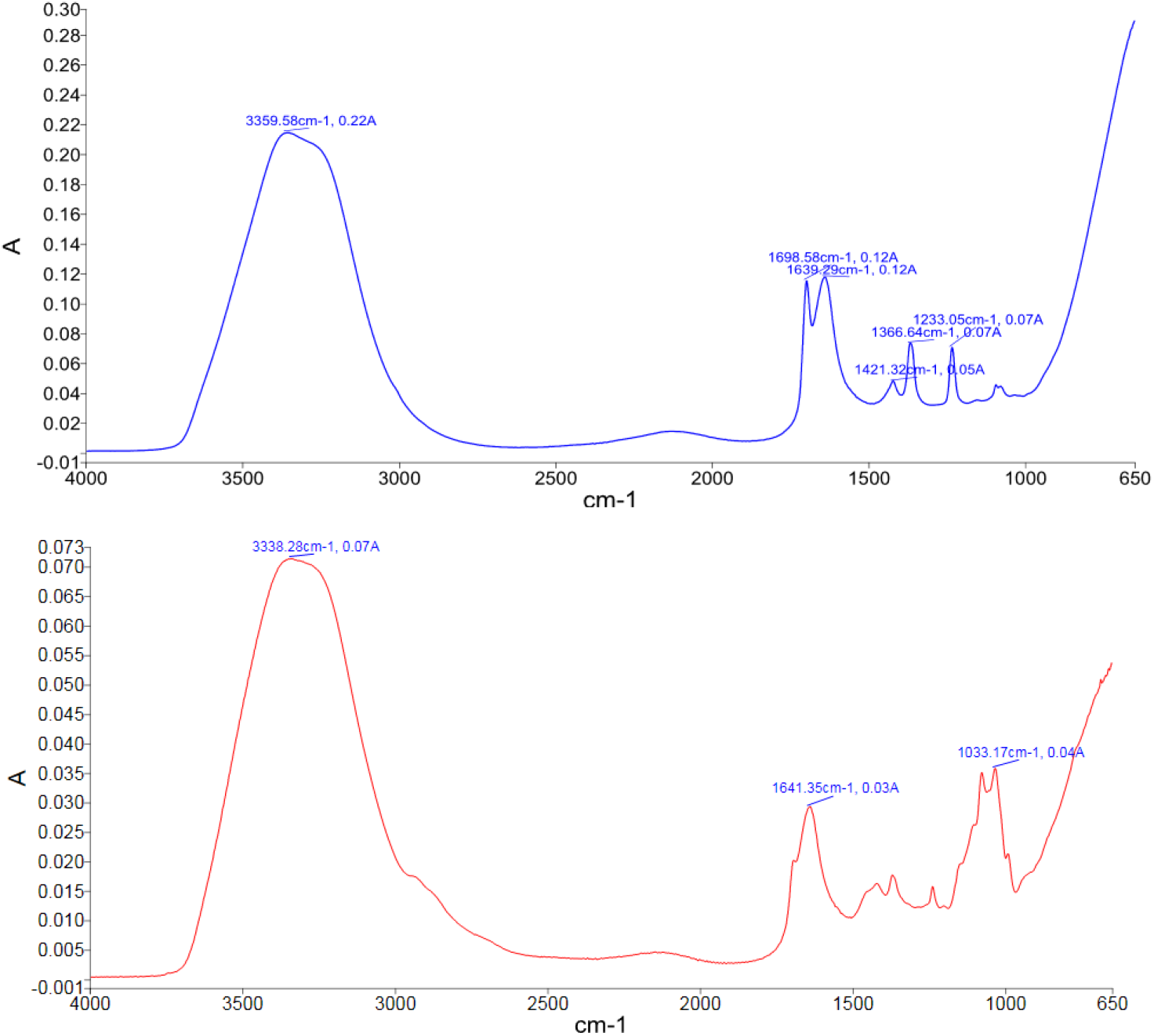
FTIR spectra of culture supernatant from control and Pb-exposed *E. coli*, showing oxalic acid–metal complexation signatures.

### Bioaccumulation Efficiency of *E. coli* for Pb and Cd Removal

The bioaccumulation efficiency of *E. coli* K-12 MG1655 to eliminate Pb and Cd in aqueous solutions was determined across varying metal concentrations, ranging from 100 ppm to 1000 ppm (Figure 3). The results demonstrate a highly effective bioaccumulation capacity of *E. coli* for Pb, Cd, and their combination. *E. coli* at 100 ppm had a Pb removal efficiency of 99.48% and Cd removal efficiency of 81.44%. Increasing concentration to 200 ppm significantly increased Cd removal to 93.76% (p = 0.02). At 500ppm, the efficiencies increased to 99.80% for Pb and 96.96% for Cd. *E. coli* was at its optimum at the highest concentration of 1000 ppm, with a 99.94% removal of Pb and 97.77% of Cd. The overall removal efficiency was 98.19% for 1000ppm Pb + Cd. These results highlight the great potential of *E. coli* in bioaccumulating Pb and Cd, with the removal of Pb always higher than the removal of Cd at all concentrations.

**Figure 3.**
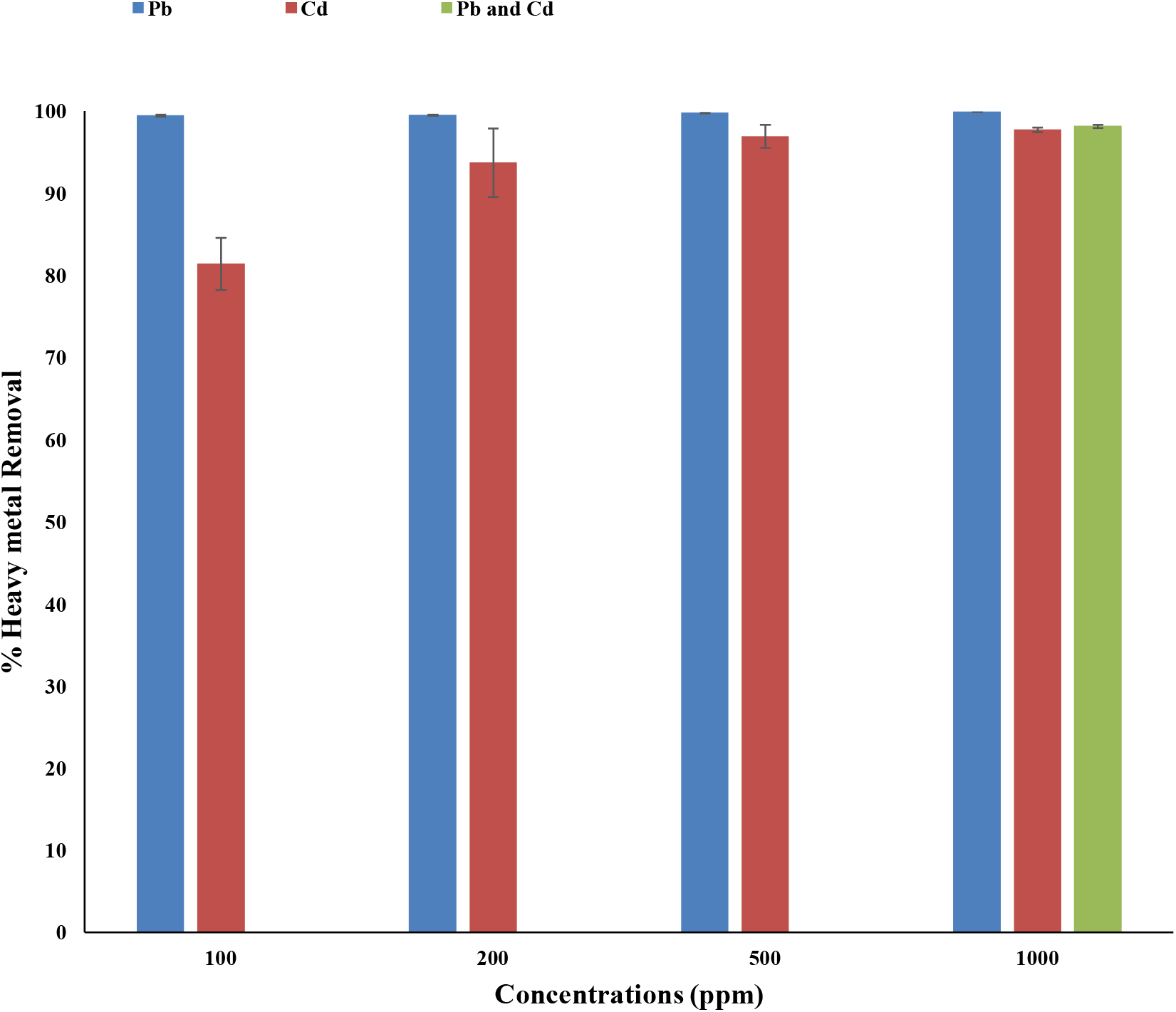
Bioaccumulation Efficiency of *E. coli* for Pb and Cd. Mean = ± SD; N = 3

### Desorption Efficiency of Pb and Cd from *E. coli* Biomass Using HCl and HNO_**3**_

The efficiency of desorption with *E. coli* K-12 MG1655 biomass was assessed at varying concentrations (0.01 M and 0.1 M) of HCl and HNO_3_. In the case of Pb, the maximum desorption efficiency (98.5%) was achieved using 0.1M HNO_3_ and then 95.46% for 0.1M HCl. Weaker solutions (0.01M HCl and 0.01M HNO_3_) gave lower recoveries of 56.78% and 71.11% respectively (Figure 4). Cd followed the same pattern: desorption efficiency was 91.5% and 85.81% with 0.1M HNO_3_ and 0.1M HCl respectively. Lower concentrations (0.01M HCl and 0.01M HNO_3_) had much lower recoveries of 51.78% and 64.61% (Figure 5). These findings suggest that 0.1 M HNO_3_ is the most effective desorbing agent of both Pb and Cd, and the concentration and type of acid used are critical in improving the desorption of heavy metals from *E. coli* biomass.

**Figure 4.**
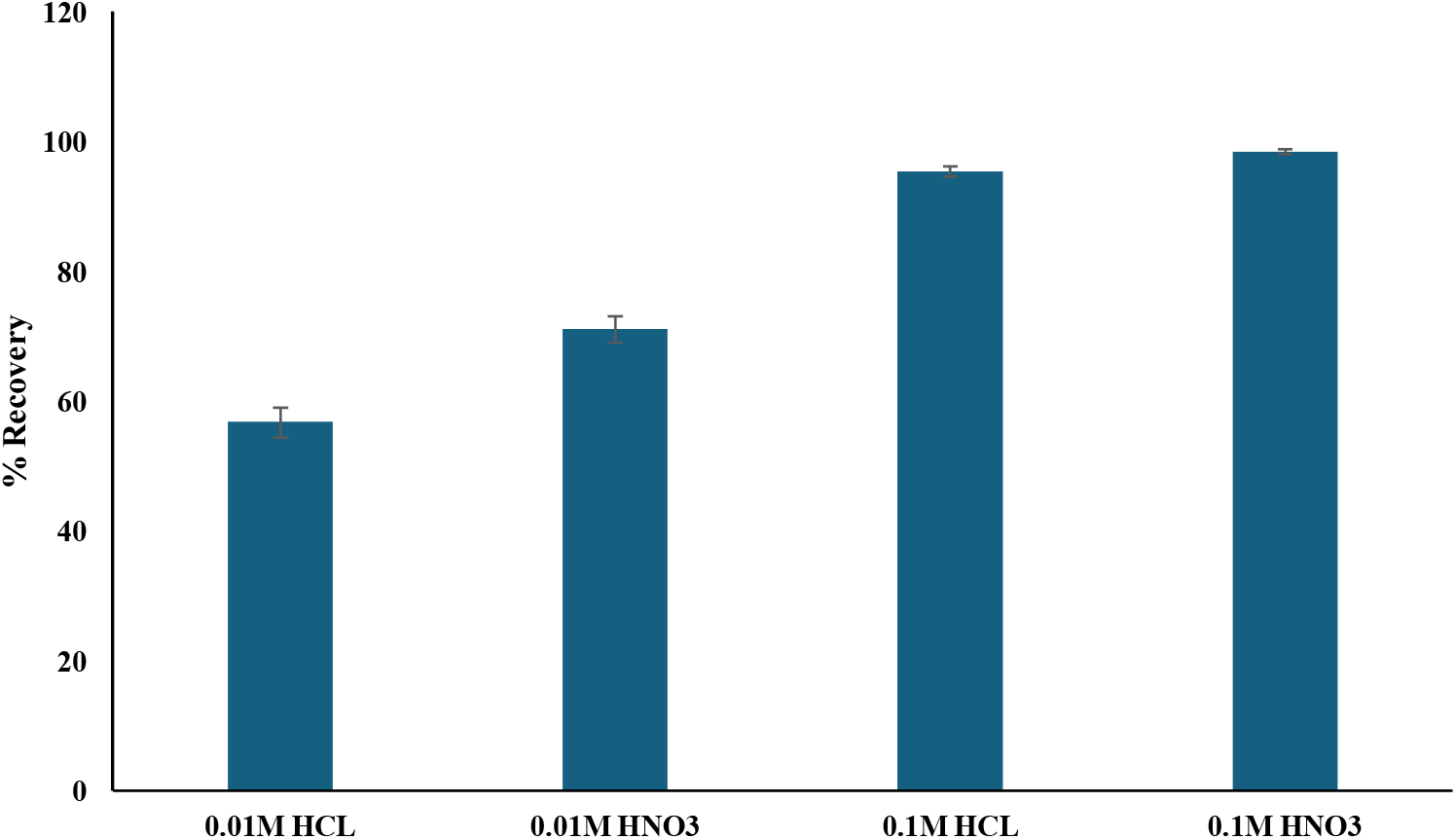
Desorption Efficiency of Pb from *E. coli* Biomass Using Different Concentrations of HCl and HNO_3_. Mean = ± SD; N = 3

**Figure 5.**
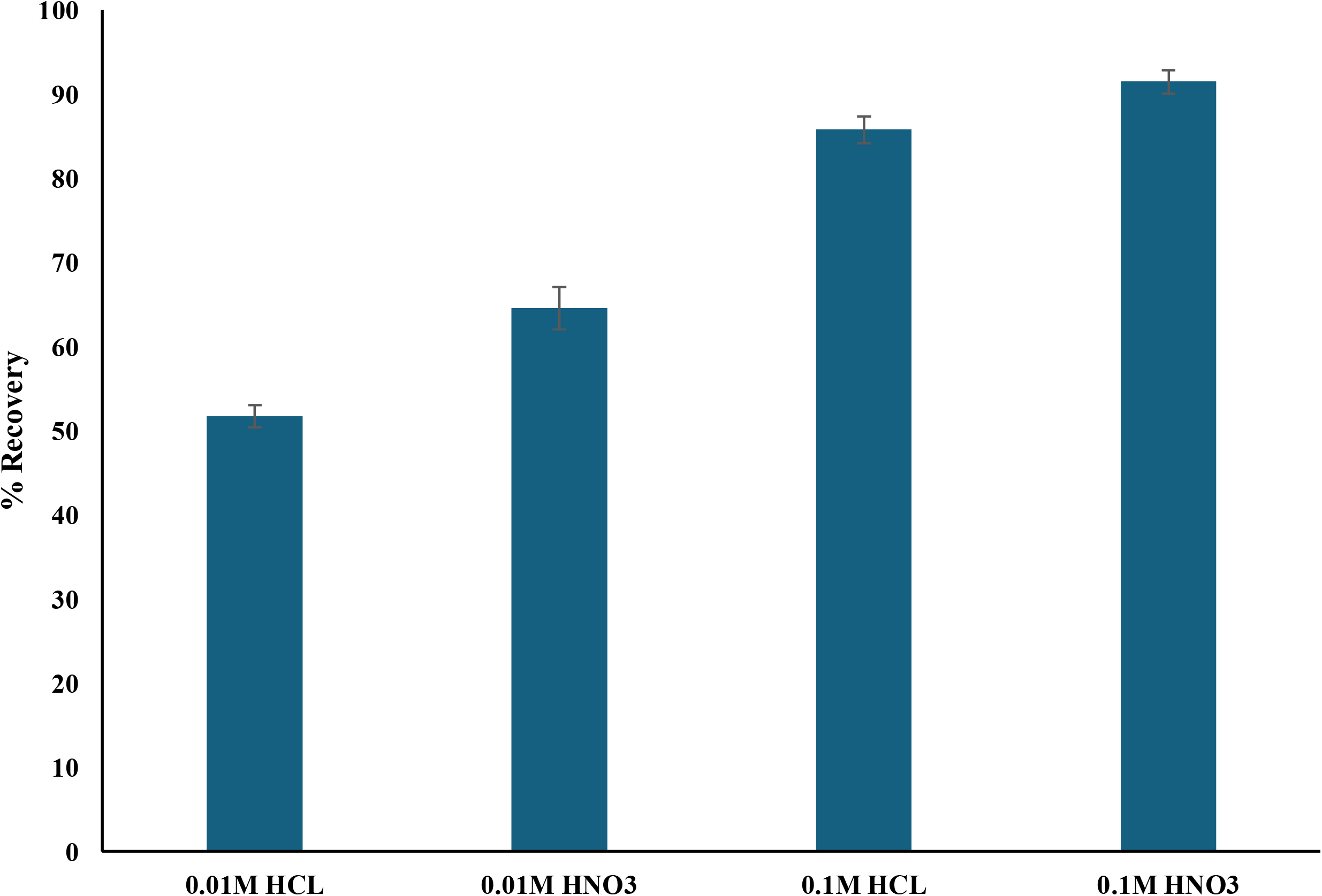
Desorption Efficiency of Cd from *E. coli* Biomass Using Different Concentrations of HCl and HNO_3_. Mean = ± SD; N = 3

### Desorption of Pb and Cd from *E. coli* Cells Using Ohmic Heating

The desorption efficiency of Pb and Cd from *E. coli* K-12 MG1655 cells using Ohmic heating was assessed by applying a controlled electrical current to cell suspensions (Figure 6). The recovery rates were measured after 10 minutes of treatment: Cd had a desorption efficiency of 45.83% and Pb had a desorption efficiency of 45.38%. These findings suggest that desorption of the heavy metals in the cells of the *E. coli* bacteria can be obtained using ohmic heating, which makes this method a possible alternative to the conventional acid-based desorption procedures.

**Figure 6.**
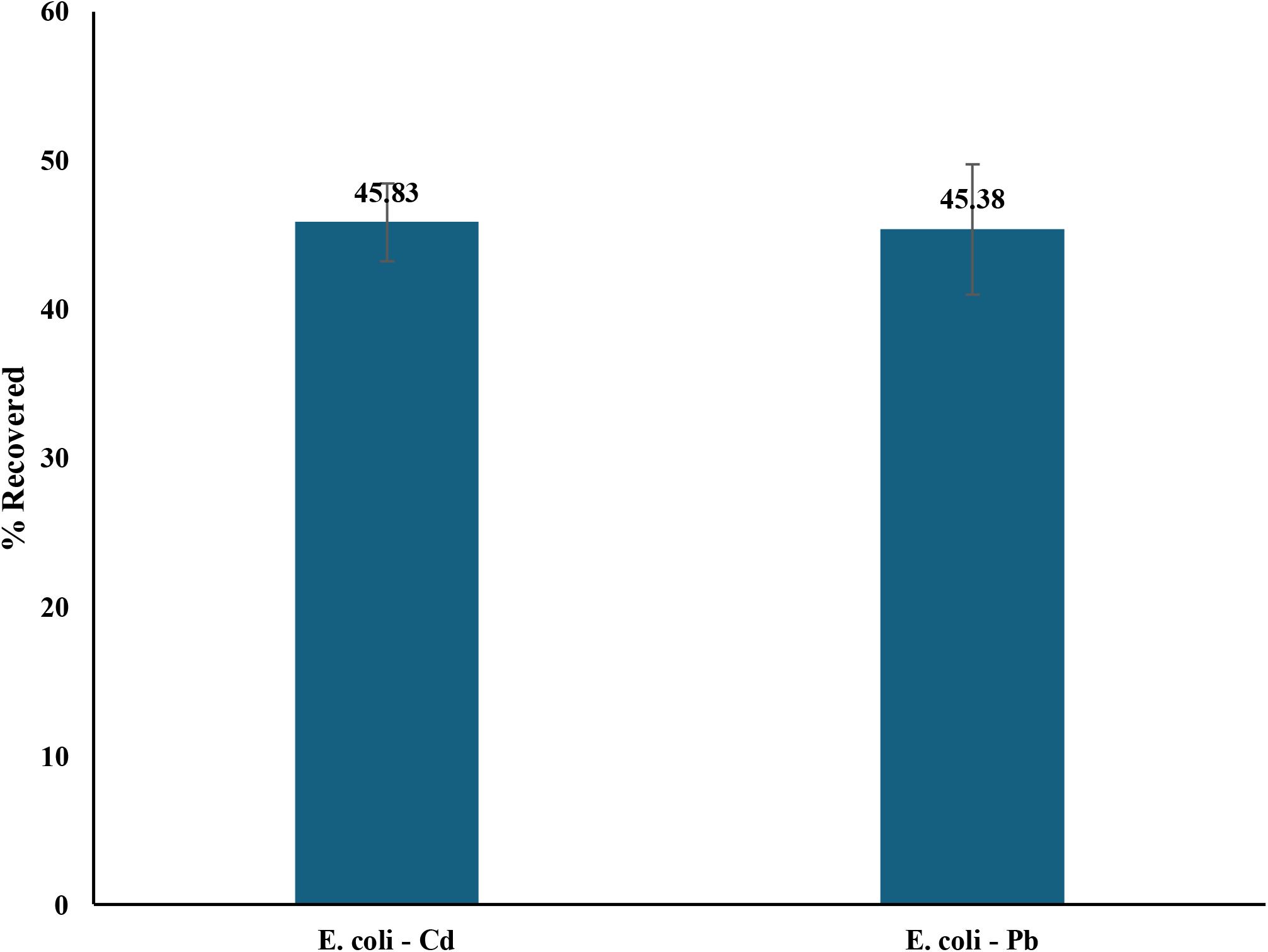
Desorption Efficiency of Pb and Cd from *E. coli* Biomass Using Ohmic Heating. Mean = ± SD; N = 3

## Discussion

The present study demonstrates that *E. coli* K-12 MG1655 adaptation to heavy-metal stress is a multifaceted biochemical and physiological process, and the production of oxalic acid is a significant detoxification pathway. Oxalic acid acts as a chemical defence molecule capable of forming stable, insoluble complexes with divalent metal ions such as Pb^2+^ and Cd^2+^. Chromatographic results indicate that the production of oxalic acid in Pb-exposed setup increased, whereas Cd exposure produces moderate increase, which confirms that the production of organic-acids is metal-specific and stress-responsive [19, 20]. Organic acids have been extensively reported in detoxification by microbial taxa; oxalate is produced by species including *Aspergillus niger* and *Bacillus subtilis* to precipitate toxic metals in inert minerals [21, 22]. This comparative evidence indicates that oxalic acid secreted by the culture of *E. coli* stabilises metal ions through the formation of the oxalate complex, and, therefore, decreases oxidative stress and preserves intracellular structures. In real world systems, microorganisms that generate organic acids are often enriched in soils contaminated with metals where the precipitation of metal-oxalate is a critical component of the natural detoxification process [23, 24]. As a result, the increased production of oxalic acid at Pb exposure does not only substantiate a biochemical stress response but also places *E. coli* in the context of microbial strategies of adapting to toxic environments in general, and the argument that metabolite-based detoxification is central to microbial metal tolerance.

FTIR analysis of the Pb-treated supernatant revealed a marked attenuation of major organic functional-group bands relative to the control, together with the emergence of a distinct absorption at ∼1033 cm^−1^. This spectral pattern is consistent with extracellular oxalic acid reacting with Pb^2+^ to form Pb–oxalate species, which consume free carboxylate groups and generate characteristic C–O–Pb vibrational modes in the 1030–1040 cm^−1^ region [25, 26]. Vibrational reference data further indicate that oxalate and metal–oxalate salts exhibit strong absorptions near 1310–1320 cm^−1^ and ∼1030 cm^−1^, supporting assignment of the new peak to oxalate–Pb coordination [27, 28]. Biologically, the formation of such complexes aligns with established microbial detoxification strategies, as many bacteria and fungi secrete oxalic acid under metal stress and immobilize metals through precipitation of insoluble oxalate phases [23, 26]. Collectively, these spectral changes provide strong evidence that extracellular oxalic acid produced by *E. coli* actively mediates Pb complexation in the supernatant, contributing to Pb immobilization via formation of Pb–oxalate species.

Building on this biochemical observation, a crucial point concerns the mechanistic link between oxalic acid secretion and subsequent metal uptake The exceptionally high bioaccumulation efficiencies (99.94% of Pb and 97.77% of Cd at 1000ppm) indicate that *E. coli* has an outstanding ability to bind and retain metal ions. This is in line with known facts that metal biosorption occurs through a primary interface with the bacterial cell envelope, which contains electronegative functional groups in abundance [29, 30]. When considered together with the oxalic acid profiles, these findings imply that oxalate does not merely detoxify but may enhance metal binding by altering local chemical environments. Organic acids can acidify microenvironments, increase ligand availability, and form bridging complexes that stabilise metal species at the cell surface. The potential role of microbial low-molecular-weight organic acids in metal complexation have been reported [31]. Thus, while oxalic acid formation primarily reduces toxicity, it can also lead to metal uptake by enhancing the probability of complexation or nucleation processes at the cell envelope. This aligns with the studies which report that detoxification and uptake interact in an integrated manner with extracellular metabolites determining the effectiveness of bioaccumulation [32, 33].

Additional evidence of this argument includes structural and biochemical findings of previous studies, including FTIR-detected protein, polysaccharide and carboxyl group vibration variations [34] and SEM-observed changes in cell-surface under metal stress [35]. The latter changes are consistent with previous studies that showed that metabolic stress induces EPS reorganization and exposure to functional groups, hence, altering metal affinity [36]. These data provide conceptual evidence that detoxification metabolites and cell-surface chemistry operate in tandem to optimise uptake. The present results therefore contribute not only experimental confirmation of high uptake rates but mechanistic depth explaining how metabolite-augmented biosorption could occur in *E. coli* under Pb and Cd stress.

This study further highlights exceptional bioaccumulation efficiency displayed by *E. coli* relative to other microbial systems. Removal rates of Pb and Cd of up to 99% and 98% respectively put *E. coli* at a very high-performance level of known biosorptive microorganisms. As an example, fungi like *Aspergillus* usually exhibit Pb removal efficiencies of 80 - 90% at similar concentrations [37] whereas many bacterial systems, including *Pseudomonas aeruginosa* or *Bacillus subtilis*, usually exhibit Cd removal of 60 – 85% under similar conditions [38, 39]. Even some of the highly touted biosorbents such as *Saccharomyces cerevisiae* rarely reach a 90% removal of Pb in single-metal systems [40]. The relatively good performance of *E. coli* in the current study can be explained by its physiological plasticity, rapid growth, and strong responses to stress, which has long been used as a model organism in metabolic studies [41, 42]. Importantly, in mixed-metal systems, where competitive inhibition often suppresses biosorption capacity, *E. coli* still retained >98% removal efficiency, indicating strong resilience to competing ions. Given that environmental contamination rarely involves single-metal exposure, this robustness under multi-metal stress enhances the ecological validity of *E. coli* as a bioremediation agent. Linking back to the aim of this study, these results strengthen the hypothesis that *E. coli* is metabolically capable as well as operationally beneficial in comparison to more specialised microbial systems.

Another critical component of this study centres on biorecovery, an aspect mostly overlooked in the conventional biosorption research. The results of desorption efficiency indicate that there is a distinct order of performance in the release of the metals in terms of recovery: 0.1 M HNO_3_ used was the most effective in releasing the metals (98.5% of Pb and 91.5% of Cd), then 0.1 M HCl, and lower concentrations of the two acids were significantly (p = 0.013) less effective. These results correspond to the chemical behaviour of metal-biomass interactions with stronger acids having higher proton-exchange capacity and destabilising metal-ligand complexes more efficiently [43, 44]. The high performance of nitric acid may be explained by its oxidative nature which helps to break down biomolecular matrices and release metal ions. Nevertheless, acid desorption is efficient; however, it increases sustainability concerns. Acid use generates secondary waste streams and introduces corrosion risks, limiting industrial applicability unless recovery circuits are tightly controlled. Therefore, the inclusion of Ohmic heating is particularly significant. This physicochemical technique provides a chemical-free, energy-driven alternative, which is in line with the concept of green-technology, although its recovery efficiencies were moderate compared to acid desorption. Ohmic heating has also been suggested as a possible technique to disinfect wastewater, and the method is already used in food processing, where it has the benefit of being easy to manage and scale [18, 45, 46]. In a circular bioeconomy model, which is defined by recovery, reuse, and the minimum amount of waste generation, this strategy has a significant potential in case of additional optimisation. As a result, biorecovery findings do not only provide quantitative data, but also conceptual information on how microbial bioremediation can be integrated into sustainable industrial processes.

The final point of analysis concerns the environmental significance and translational potential of the integrated detoxification–uptake–recovery model demonstrated in this study. Heavy-metal contamination continues to be a critical ecological and population-health problem globally [47]. Traditional methods of treatment, which include precipitation, ion exchange and electrochemical remediation, are expensive and have low efficiency at dilute concentrations [6, 48]. The current results suggest that microbial systems would be a viable alternative: not only does the system of microbes detoxify and accumulate metals with a high level of efficiency but also allows the recovery of the metals. This is a twofold ability that is necessary in the development of circular remediation systems that would be able to attain both environmental purification and resource recovery. A microbial system with the ability to simultaneously sequester and recover could greatly lower the operational costs and reduce the amount of chemical input in industrial applications (i.e. battery recycling, mining effluent treatment, and electronic-waste processing). In addition, the metabolic interdependence between oxalic-acid synthesis and metal uptake is observed to give mechanistic roadmap to strain engineering in the future. By optimising the microbial recovery pathways further, membrane engineering, or transporter overexpression may be a possibility through increased organic-acid secretion. Real-world analogues exist in wastewater bioreactors and bioleaching operations, where microbial consortia are engineered to improve metal capture and yield. Therefore, the current work provides evidence of the feasibility of the concept under consideration, as well as outlines some of the future avenues of practical implementation.

Taken together, our results move beyond conventional biosorption reports that treat detoxification, removal and recovery as disconnected operations. By integrating oxalate-mediated detoxification, high-capacity bioremoval and subsequent metal recovery within a single *E. coli* K-12 MG1655 platform, we demonstrate a coherent workflow for managing Pb and Cd at high loadings. The deployment of ohmic heating as a regeneration and recovery step for metal-loaded bacterial biomass is particularly notable, offering a low-chemical, energy-efficient alternative to conventional acid stripping that, to our knowledge, has not been systematically explored for Pb/Cd systems. This integrated perspective underscores the potential of *E. coli*–based processes as flexible, regenerable tools for heavy metal management.

## Conclusion

This study demonstrates that *E. coli* K-12 MG1655 uses a metabolically coordinated response to tolerate and process heavy-metal stress, and that the production of oxalic-acid can be considered as a major part of its detoxification program. The organism produced detectable levels of oxalic acid under Pb and Cd exposure, with Pb eliciting the strongest response, underscoring the role of oxalate in mitigating toxicity through complexation and precipitation of metal ions. This biochemical adaptation was highly consistent with the extremely high bioaccumulation potential of the organism, where both metals and nearly full uptake of Pb were removed at the highest levels. Such performance positions *E. coli* K-12 MG1655 among the most efficient microbial metal bioaccumulators. Given this performance, the system shows strong potential for scale-up using industrial bioreactor configurations, including packed-bed or fluidised-bed systems, and could be integrated into existing wastewater treatment lines as a modular metal-removal unit

In parallel, the study established viable recovery pathways capable of reclaiming the sequestered metals from the biomass. The optimum recovery was obtained using 0.1M HNO_3_ where the maximum percentage of Pb and Cd was 98.5% and 91.5% respectively, which means that biosorbents can be chemically regenerated. Though the recovery rates of ohmic heating were relatively low, its chemical-free and scalable characteristics make it preferable to use as an environmental-friendly solution, which can be optimised to be used in industry.

In sum, these findings demonstrate a complete and functional microbial remediation recovery cycle all the detoxification, sequestration, and metal recovery are mechanistically research paper makes *E. coli* K-12 MG1655 a viable candidate in closed-loop bioremediation systems that involve detoxification, sequestration, and recovery of heavy metals.

## Author Contributions

**N.D.N:** Writing – review & editing, Writing – original draft, Methodology, Investigation, Formal analysis, Data curation, Conceptualization. **C.U.A:** Writing – review & editing, Supervision, Conceptualization. **T.M:** Writing – review & editing, Supervision, Conceptualization. **H.O:** Writing – review & editing, Supervision, Conceptualization.

